# Spindle checkpoint signalling in anaphase is prevented by KNL1 release from kinetochores

**DOI:** 10.1101/2023.05.10.540295

**Authors:** Iona Lim-Manley, Ulrike Gruneberg

**Author notes:** Current address: Organelle Dynamics Laboratory, The Francis Crick Institute, London, UK. corresponding author Please address all correspondence to: Ulrike Gruneberg.

## Abstract

CDK1-cyclin B1 kinase is the main driver of mitosis and initiates the morphological changes that characterise mitosis, including mitotic spindle assembly and formation of the outer kinetochore. CDK1-cyclin B1 activity is also critically required for spindle assembly checkpoint (SAC) signalling during mitosis. In particular, CDK1-cyclin B1 promotes the targeting of the principal spindle checkpoint kinase MPS1 to kinetochores, leading to the recruitment of SAC proteins to the outer kinetochore scaffold protein KNL1 and initiation of checkpoint signalling. However, cells expressing kinetochore-tethered MPS1 still require CDK1 activity for SAC signalling, suggesting that CDK1 plays both MPS1-dependent and -independent roles in regulating the SAC. Here we show that the latter is due to CDK1-mediated kinetochore recruitment of KNL1, which is reversed by the PP1 phosphatase at the metaphase-to-anaphase transition. Our findings explain the abrupt and irreversible termination of spindle checkpoint signalling in anaphase, since the drop of CDK1 activity means both MPS1 and the spindle checkpoint scaffold KNL1 are lost from kinetochores.

**Summary:** Lim-Manley and Gruneberg investigate MPS1-independent roles of CDK1 in spindle checkpoint signalling. They reveal how PP1 activity following CDK1 inactivation results in the rapid removal of KNL1 from kinetochores at anaphase onset, contributing to prompt spindle checkpoint silencing.

## Introduction

The accurate segregation of chromosomes in mitosis is essential for the faithful inheritance of genetic material. The fidelity of chromosome segregation is monitored by the spindle assembly checkpoint (SAC), an important cellular pathway that ensures that all chromosomes have formed stable attachments to microtubules via their kinetochores before chromosome segregation ensues (Lara-Gonzalez et al., 2012; Musacchio, 2015). If missing or erroneous microtubule-kinetochore attachments are detected, progression into anaphase is prevented until the problem has been rectified. MPS1, the principal SAC organizer, is recruited to the NDC80 complex of unattached or incorrectly attached kinetochores and from there initiates spindle checkpoint signalling by phosphorylating the outer kinetochore scaffold protein KNL1 on recurrent Met-Glu-Leu-Thr (MELT) motifs (Ji et al., 2015; London et al., 2012; Nijenhuis et al., 2013; Pachis and Kops, 2018; Shepperd et al., 2012; Yamagishi et al., 2012). These phosphorylated sites generate landing pads for SAC proteins of the BUB family, which in turn recruit the MAD1 and MAD2 proteins, initiating the catalytic formation of the mitotic checkpoint complex (MCC), inhibition of the anaphase promoting complex/cyclosome (APC/C) by the MCC, and consequent cell cycle arrest until all chromosomes have been securely attached to microtubules (Musacchio, 2015). Once this has been achieved, the spindle assembly checkpoint is silenced and cells are able to exit mitosis.

A key cellular structure during chromosome segregation is the kinetochore, the multi-layered proteinaceous structure, which assembles on the centromeric chromatin at the beginning of mitosis. In particular, the KMN network of the outer kinetochore, consisting of the KNL1-ZWINT complex, the MIS12 complex, made up of the MIS12, DSN1, NSL1 and NNF1 proteins, and the NDC80 complex containing NDC80, NUF2, SPC24 and SPC25, fulfils two critical functions during chromosome segregation. On the one hand it provides the physical connection between the chromosomes and the microtubules, but on the other hand it also acts as the assembly platform for the SAC (Cheeseman et al., 2006; Musacchio and Desai, 2017). Continued spindle assembly checkpoint signalling throughout prometaphase is very important for achieving bi-orientation and hence accurate chromosome segregation, but at the same time it is paramount that the spindle checkpoint is silenced promptly in anaphase and not re-activated so chromosome segregation, once started, is not abrogated. SAC responsiveness is therefore limited to a specific time window during mitosis and requires CDK1 activity for both the initiation and maintenance of the SAC signal (D’Angiolella et al., 2003; Rattani et al., 2014; Vazquez-Novelle et al., 2014). One key target of CDK1-cyclin B1 in the spindle checkpoint is MPS1, which is phosphorylated in the kinetochore binding domain by CDK1, promoting kinetochore binding (Hayward et al., 2019). SAC responsiveness during mitosis is hence only possible as long as MPS1 is phosphorylated (Hayward et al., 2019). Apart from MPS1, BUB1 has been shown to be another important CDK1 target within the SAC, as it is sequentially phosphorylated by CDK1 and MPS1 to aid MAD1 binding (Ji et al., 2017; Qian et al., 2017; Zhang et al., 2017). Intriguingly, cyclin B1 itself localizes to checkpoint active kinetochores by binding to MAD1, potentially aiding spindle checkpoint signalling at low CDK1-cyclin B1 activity through a positive feedback loop (Alfonso-Perez et al., 2019; Allan et al., 2020; Jackman et al., 2020). On the kinetochore side, CENP-T is a known CDK1 target which needs to be phosphorylated for KNL1/MIS12 complex and NDC80 complex recruitment in late G2/early mitosis (Gascoigne and Cheeseman, 2013; Huis In ’t Veld et al., 2016; Rago et al., 2015).

Timely dephosphorylation of these CDK1 substrates is critical for the correct temporal orchestration of SAC silencing and progression through mitosis and anaphase, and is mainly carried out by phosphatases of the PP2A and PP1 families, in particular PP2A-B55 and PP1αγ (Bancroft et al., 2020; Cundell et al., 2013; Cundell et al., 2016; Gascoigne and Cheeseman, 2013; Hayward et al., 2019; Holder et al., 2020; Schmitz et al., 2010). Of these, PP2A-B55 opposes the CDK1 phosphorylation of MPS1 to terminate the SAC permissive period by preventing further MPS1 kinetochore recruitment (Hayward et al., 2019).

The identification of the phosphatases opposing specific mitotic phosphorylation events is an important ongoing task, and is tightly linked to the discovery of further CDK1 targets required for spindle checkpoint signalling. To identify such additional CDK1 substrates, here we used cells expressing a MIS12-MPS1 fusion protein which tethers MPS1 to kinetochores and thus circumvents the need for CDK1 activity for MPS1 kinetochore targeting. This enabled us to investigate MPS1-independent roles of CDK1-cyclin B1 in spindle checkpoint signalling. Our analysis revealed a specific role for CDK1 in the recruitment of the KNL1 spindle checkpoint scaffold critical for the assembly of spindle checkpoint proteins. This event is opposed by PP1 in anaphase, contributing to the timely and irreversible silencing of the spindle assembly checkpoint.

## Results and Discussion

### Kinetochore recruitment of checkpoint proteins requires CDK1, independently of MPS1

We have previously shown that the requirement for CDK1 for MPS1 localisation can be circumvented by fusing MPS1 to the outer kinetochore protein MIS12 (Hayward et al., 2019; Jelluma et al., 2010). We therefore used this approach to test whether checkpoint signalling requires CDK1 activity for substrates other than MPS1. We generated HeLa Flp-In T-REx cells expressing the GFP-MIS12-MPS1 fusion protein in the absence of endogenous MPS1 and monitored the kinetochore localisation of key spindle checkpoint proteins (BUB1, BUBR1, MAD1) in the presence or absence of a short, 6 min treatment with the CDK1 inhibitor flavopiridol (CDKi) (Losiewicz et al., 1994) (Figure 1). To assess the requirement for CDK1 activity in either the establishment or maintenance of the checkpoint, we used two different assays. To analyse spindle checkpoint re-activation, we arrested cells in metaphase using the proteasome inhibitor MG132, followed by a brief nocodazole treatment to induce mitotic spindle defects (Hayward et al., 2019; Vleugel et al., 2015) (Figure 1A). Under these conditions, cyclin B1 and securin degradation were prevented due to the presence of MG132, allowing us to study the effect of CDK1 substrate dephosphorylation upon CDK1 inhibition in isolation. Despite the presence of the MIS12-MPS1 fusion protein, treatment with CDKi resulted in the absence of BUB1, BUBR1 and MAD1 from kinetochores, suggesting a CDK1-dependent step in their kinetochore recruitment in addition to the known requirement for MPS1 (Figure 1B, C). To assess the importance of CDK1 activity for the maintenance of an already established spindle checkpoint, we arrested cells for 2 hrs in the presence of nocodazole, and then exposed them to CDKi for 6 min in the presence of the proteasome inhibitor MG132 to prevent mitotic exit (Figure 1D). In this situation, we observed that CDKi exposure also resulted in the loss of BUB1 and BUBR1 from kinetochores, but MAD1 was retained (Figure 1E and F). This difference in MAD1 behaviour is most likely due to both BUB1 and the Rod-Zw10-Zwilch (RZZ) complex contributing to MAD1 localization, with BUB1 acting as the initial, CDK1 inhibition-sensitive, MAD1 receptor, and RZZ then taking over during prolonged spindle checkpoint arrest as the main MAD1 interaction partner which is only sensitive to CDKi after a longer 20 min treatment (Rodriguez-Rodriguez et al., 2018; Sacristan et al., 2018; Silio et al., 2015). Since we were mainly interested in understanding the role of CDK1 in the initiation of SAC signalling, we decided to focus on the SAC re-activation assay for further experiments.

**Figure 1.**
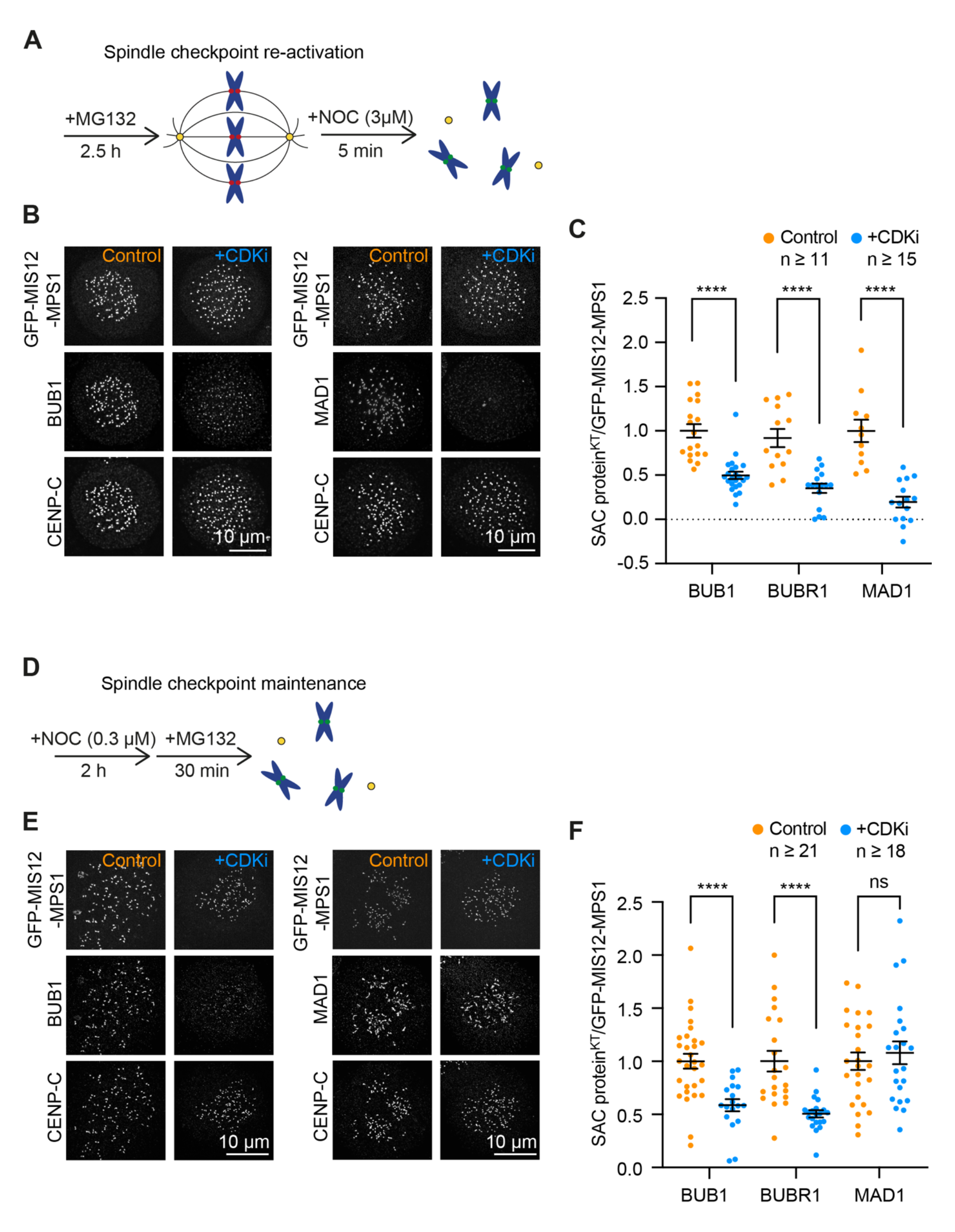
Tethering MPS1 to kinetochores is insufficient for spindle checkpoint signalling. (**A**) Schematic of the “checkpoint re-activation assay”. (**B**) HeLa Flp-In/T-REx GFP-MIS12-MPS1 depleted of endogenous MPS1 were arrested for 2.5 h with 20 μM MG132, then treated with 3 μM nocodazole for 5 min, fixed with PTEMF, and immunostained as indicated. Where indicated, 5 μM flavopiridol was added 1 min before nocodazole treatment (+CDKi). (**C**) Graph shows mean cell intensities of kinetochore BUB1, BUBR1 and MAD1 relative to GFP-MIS12-MPS1. Bars indicate SEM. (**D**) Schematic of the “checkpoint maintenance assay”. (**E**) HeLa Flp-In/T-REx GFP-MIS12-MPS1 depleted of endogenous MPS1 were arrested for 2 h with 0.3 μM nocodazole, then treated with 20 μM MG132 for 30 min, fixed with PTEMF, and immunostained as indicated. Where indicated, 5 μM flavopiridol was added 6 min before fixation (+CDKi). (F) Graph shows mean cell intensities of kinetochore BUB1, BUBR1 and MAD1 relative to GFP-MIS12-MPS1. Bars indicate SEM.

### CDK1 regulates MPS1 localisation but not its activity

MPS1 has four reported CDK1 phosphorylation sites (S281, S436, T453 and S821) (Figure S1A), and CDK1-dependent phosphorylation of human MPS1 at S821 or Xenopus MPS1 at S283, homologous to human S281, has been suggested to be important for MPS1 kinase activity (Diril et al., 2016; Dou et al., 2011; Hayward et al., 2019; Morin et al., 2012). One potential interpretation of our findings is therefore that in addition to regulating MPS1 localization, CDK1 may be required for MPS1 activity. To test this possibility, HeLa Flp-In T-REx cells expressing versions of GFP-MIS12- MPS1 in which all four phospho-sites had been mutated to phosphorylation-deficient alanine (4A) or phospho-mimetic glutamate (4E) (Figure S1B) were used in the checkpoint re-activation assay (Figure S1C and D). Cells expressing GFP-MIS12- MPS1^4A^ recruited kinetochore BUB1 to the same level as the wild type construct indicating that phosphorylation of these sites is not necessary for MPS1 activity (Figure S1C and D). Furthermore, CDKi still resulted in strongly reduced BUB1 kinetochore localization in cells expressing phospho-mimetic GFP-MIS12-MPS1^4E^, consistent with the idea that CDK1 substrates other than MPS1 are required for spindle checkpoint protein localization (Figure S1C and D). To strengthen the conclusion that CDK1 activity is not required for MPS1 catalytic activity, wild type (WT), kinase-dead (KD), S821A and 4A versions of MPS1 were expressed as FLAG-tagged proteins in HEK- 293T cells, purified and tested for their kinase activity in in vitro kinase assays. No reduction in MPS1 auto-catalytic kinase activity was observed in the S821A or 4A mutants, supporting the view that CDK1 phosphorylation is not required for MPS1 activity (Figure S1E and F).

### Kinetochore recruitment of the checkpoint scaffold KNL1 is CDK1-sensitive

Since we observed the same CDK1-dependence for BUB1, BUBR1 and MAD1 in initiating a checkpoint signal, we suspected that an upstream factor common to all three proteins was regulated by CDK1 activity. The outer kinetochore protein KNL1 constitutes the critical scaffold for spindle checkpoint protein assembly upon MPS1 phosphorylation, and hence is a potential candidate. As it is known that assembly of the outer kinetochore is regulated by CDK1-cyclin B1 activity (Gascoigne and Cheeseman, 2013; Huis In ’t Veld et al., 2016; Rago et al., 2015), we investigated the effect of CDKi treatment on KNL1 kinetochore levels in the GFP-MIS12-MPS1 cells (Figure 2A). While the kinetochore levels of HEC1, the main microtubule binding factor at the outer kinetochore, were not affected by a brief, 6 min treatment with CDKi, KNL1 kinetochore levels were strongly reduced (Figure 2A and B) by the same treatment, indicating a differential sensitivity of the different KMN complexes to CDKi. To be able to compare the behaviour of different outer kinetochore proteins without relying on antibodies with varying affinities for their antigens, we generated HeLa Cas12a-modified cell lines expressing endogenously GFP-tagged versions of the outer kinetochore proteins KNL1, HEC1 and MIS12 (Figure S2A and B). This enabled us to directly compare the sensitivities of these proteins to CDKi after 6 min and 10 min of inhibition. As already observed with antibody staining, KNL1-GFP was highly sensitive to CDK1 inhibition, and KNL1 localization was strongly reduced after 6 min of CDKi. HEC1, in comparison, was largely unchanged, while MIS12 showed an intermediate behaviour with reduced kinetochore localization visible after 10, but not 6 min of CDKi (Figure 2C and D).

**Figure 2.**
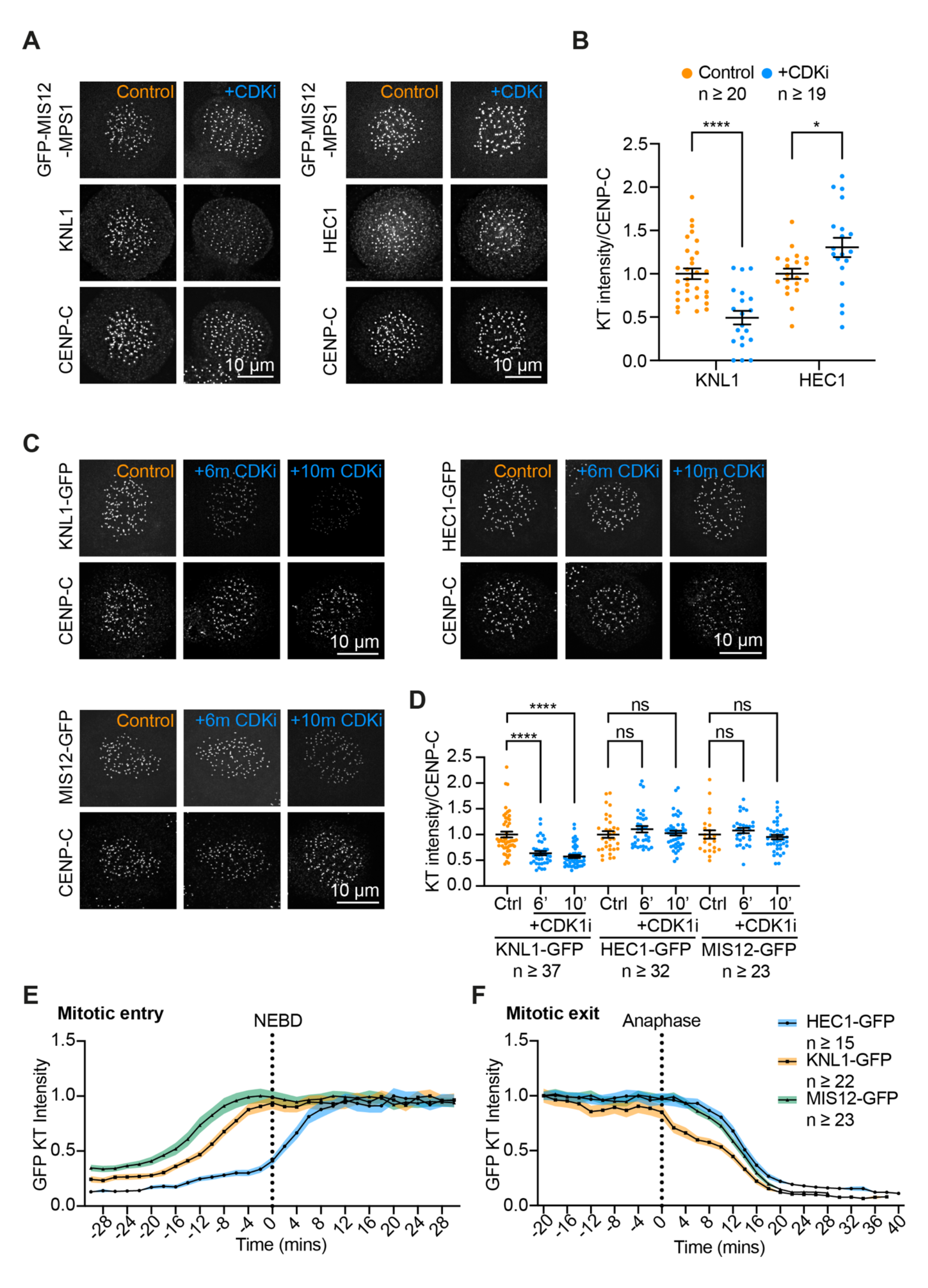
Kinetochore recruitment of KNL1 is CDK1 sensitive. (**A**) HeLa Flp-In/T-REx GFP-MIS12-MPS1 cells depleted of endogenous MPS1 were treated as in Figure 1B. (**B**) Graph shows mean cell intensities of kinetochore KNL1 and HEC1 relative to CENP-C. Bars indicate the SEM. (**C**) HeLa cells expressing endogenously tagged KNL1-GFP, HEC1-GFP or MIS12-GFP were arrested for 2.5 h with 20 μM MG132, then treated with 3 μM nocodazole for 5 min, fixed and immunostained as indicated. Where indicated, 5 μM flavopiridol was added 1 or 5 min before nocodazole treatment (+6m CDKi and +10m CDKi respectively). (**D**) Graph shows mean cell intensities of kinetochore KNL1-GFP, HEC1-GFP or MIS12-GFP relative to CENP-C. Bars indicate mean ±SEM. (**E**) HeLa cells expressing KNL1-GFP, MIS12-GFP or HEC1-GFP were synchronised for 18 h using 2 mM thymidine. Cells were incubated with 100 nM SiR-DNA and imaged 9 h after thymidine release. Graphs show mean cell intensities of kinetochore KNL1-GFP, HEC1-GFP or MIS12-GFP at 2 min intervals as a fraction of the maximum intensity. Shaded regions indicate the SEM.

Next, we compared the kinetochore assembly and disassembly kinetics of KNL1-GFP, HEC1-GFP and MIS12-GFP by live cell imaging in unperturbed cells. This analysis revealed clear differences in when KNL1, HEC1 and MIS12 are recruited to and released from kinetochores during mitotic entry and exit (Figure 2E and F, Figure S2C). In particular, KNL1 and MIS12 were recruited to kinetochores early in mitosis, before nuclear envelope breakdown, and markedly earlier than HEC1 which only appeared after nuclear envelope breakdown (Figure 2E), consistent with previous observations (Gascoigne and Cheeseman, 2013). At mitotic exit, KNL1 left the kinetochores earlier than either MIS12 or HEC1 (Figure 2E, Figure S2C). These observations, together with our CDKi experiments in Figure 2C, demonstrate that, although at the kinetochore assembly stage KNL1 and MIS12 behave essentially as a unit (Rago et al., 2015), this is not the case at mitotic exit when KNL1 is the first protein to leave the outer kinetochore upon the drop of CDK1 activity at anaphase onset, independently of MIS12.

### KNL1 recruitment to the kinetochore requires CDK1-phosphorylation of NSL1

KNL1 binds to the kinetochore via an interaction with the MIS12 complex protein NSL1, and depletion of NSL1 results in the complete loss of KNL1 from the kinetochore (Petrovic et al., 2014; Petrovic et al., 2010) (Figure S3A-C). The KNL1- NSL1 interaction involves a C-terminal segment in KNL1 encompassing amino acids 2000-2311 and a C-terminal peptide of NSL1 comprising amino acids 227-281 (Petrovic et al., 2014; Petrovic et al., 2010). Interestingly, we observed that the extended NSL1 C-terminal sequence contains a conserved consensus CDK1 phosphorylation (Thr 242) site within a threonine-rich patch in close proximity to the KNL1-interacting peptide, raising the possibility that the interaction between NSL1 and KNL1 is regulated by CDK1 phosphorylation (Figure 3A). To test this hypothesis, we first checked whether NSL1 was mitotically phosphorylated. PhosTag SDS-PAGE analysis (Kinoshita et al., 2006), which allows the visualization of phosphorylated protein species, of HeLa cell extracts from asynchronous or mitotically arrested cells showed a significant upshift of NSL1 in mitotic cells (Figure 3B). This upshift was fully reversed by the addition of λ-phosphatase to the mitotic extract, confirming that the upshift was due to phosphorylation and not other post-translational modifications (Figure 3B). In addition to this, mass-spectrometry analysis of overexpressed GFP-NSL1 purified from mitotic HEK-293T cells demonstrated that Thr 242 was phosphorylated in mitosis (Figure 3C). In line with the idea that the interaction between NSL1 and KNL1 was phosphorylation sensitive, we found that in immunoprecipitations of GFP-NSL1, the interaction with KNL1 was strongly diminished when the cells had been treated with CDKi before immunoprecipitation (Figure 3D). To analyse the consequences of phospho-deficient or phospho-mimetic mutants at the Thr 242 site for KNL1 localization, mitotically arrested HeLa Flp-In T-REx cells depleted for endogenous NSL1 and expressing either wild type GFP-NSL1 (GFP-NSL1^WT^), or GFP-NSL1 with both Thr 242 and adjacent Thr 241 mutated to either phospho-null alanine (GFP-NSL1^2TA^) or phospho-mimetic aspartate (GFP-NSL1^2TD^) in the presence or absence of CDK1 inhibitor, were fixed and stained for KNL1 (Figure 3E, Figure S3D). As an additional control, we used a version of GFP-NSL1 in which tyrosine 270, critical for the interaction with KNL1, had been mutated to alanine (GFP-NSL1^Y270A^) (Petrovic et al., 2014; Petrovic et al., 2010). GFP-NSL1 expression was comparable in all cases, indicating that the mutations did not affect NSL1 stability (Figure S3D). GFP-NSL1^2TA^ and GFP-NSL1^2TD^ targeted to kinetochores as well as GFP-NSL1^WT^, suggesting that the mutations did not affect their ability to incorporate into the MIS12 complex (Figure 3E). GFP-NSL1^Y270A^ kinetochore localization, however, was reduced, in line with the idea that this mutation disturbed kinetochore assembly. Furthermore, expression of GFP-NSL1^Y270A^ nearly abrogated KNL1 kinetochore binding, confirming the published data (Petrovic et al., 2014). Interestingly, in GFP-NSL1^2TA^ cells, KNL1 kinetochore recruitment was significantly reduced, but not completely abolished, whereas expression of GFP-NSL1^2TD^ re-established KNL1 localization to wild type levels (Figure 3E and F). Using immunoprecipitations of the different forms of GFP-NSL1, a similar pattern was observed biochemically, with a loss of both KNL1 and HEC1 pull-down with GFP-NSL1^Y270A^, reduced KNL1 pull-down with GFP-NSL1^2TA^ and near restoration of KNL1 binding for GFP-NSL1^2TD^ (Figure 3G and H). Taken together, our data indicate that the kinetochore localization of KNL1 is promoted by CDK1 phosphorylation of its kinetochore binding partner, NSL1.

**Figure 3.**
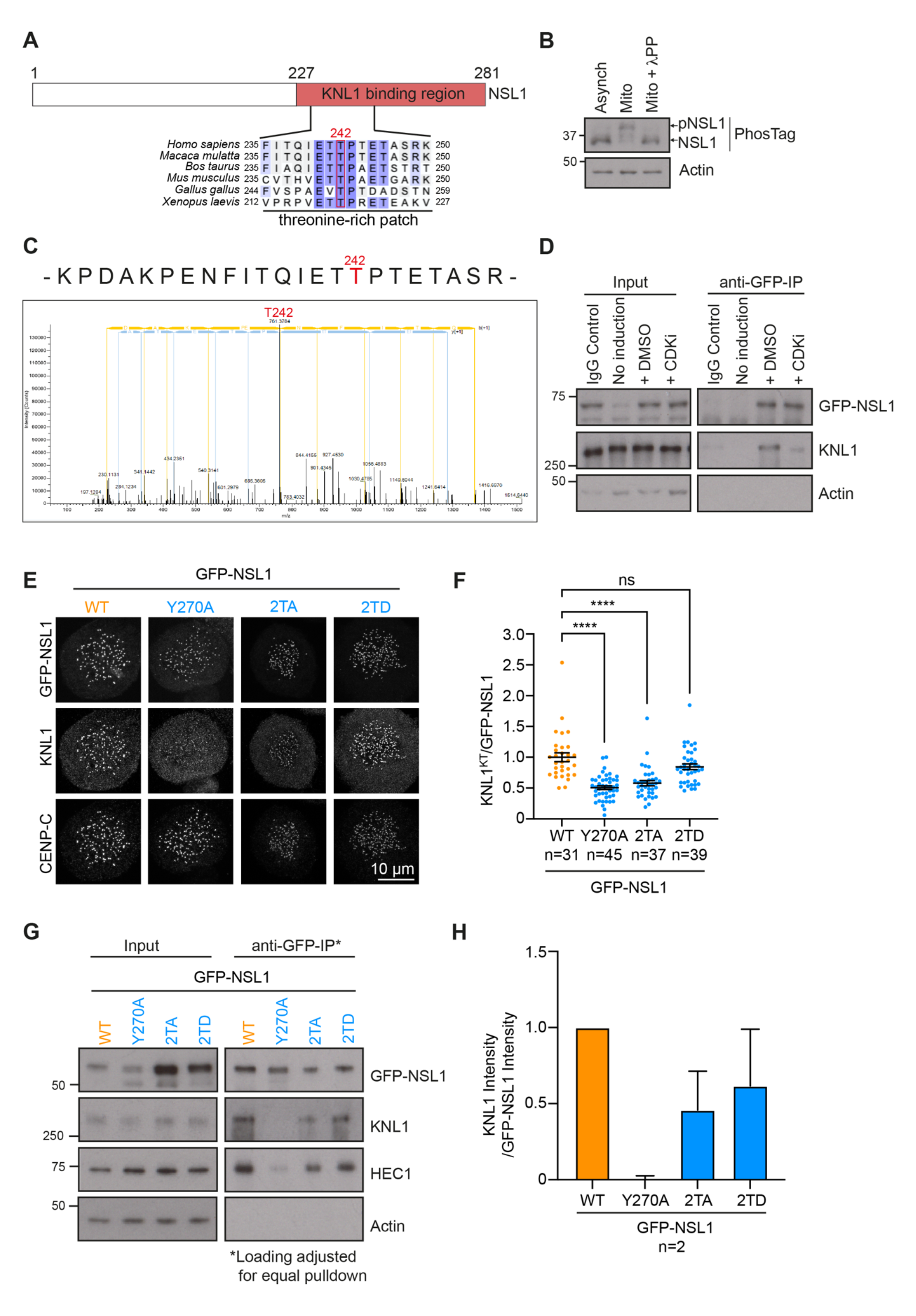
KNL1 kinetochore recruitment requires CDK1-phosphorylation of NSL1. (**A**) Schematic of domain organization of NSL1. Essential residue Y270 and proposed CDK1 phosphorylation site required for KNL1 interaction are indicated. CDK1 consensus site (T242) is shown in red. (**B**) Western blot of asynchronous and mitotic HeLa cell lysates. Mitotic lysates were treated with λ-phosphatase where indicated. Samples were run on either standard or PhosTag SDS-PAGE gels as indicated. (**C**) Graph shows m/z data from mass spectrometry analysis of GFP-NSL1 T242 phosphorylation. Peptide detected in phosphoproteomic analysis is shown above. (**D**) HeLa Flp-In/T-REx GFP-NSL1 cells were arrested for 18 h with 0.3 μM nocodazole. Where indicated, 5 μM flavopiridol was added 10 min before cells were collected and lysed. GFP-NSL1 was immunoprecipitated (IP) with anti-GFP antibodies. Immunoprecipitated protein complexes were analysed by western blotting. 0.5% of Input and 5% of IP was loaded on gels. (**E**) HeLa Flp-In/T-REx GFP-NSL1 cells depleted of endogenous NSL1 were arrested for 2.5 h with 20 μM MG132, then treated with 3 μM nocodazole for 5 min, fixed and immunostained as indicated. (**F**) Graph shows mean cell intensities of kinetochore KNL1 relative to GFP-NSL1. Bars indicate the SEM. (**G**) HeLa Flp-In/T-REx GFP-NSL1 cells arrested for 18 h with 0.3 μM nocodazole were collected and lysed. GFP-NSL1 was immunoprecipitated (IP) with anti-GFP antibodies. Immunoprecipitated protein complexes were analysed by western blotting. 0.5% of Input was loaded on gels, IP loading was adjusted for equal GFP-NSL1 pulldown. (H) Graph shows mean KNL1 signal relative to GFP-NSL1 pulldown. Bars indicate S.D.

### PP1 promotes timely release of KNL1 from kinetochores in anaphase

Our results so far indicated that in anaphase KNL1 is lost from kinetochores earlier than the other outer kinetochore proteins. This would imply differential phosphatase action, preferentially removing CDK1 phosphorylations promoting KNL1 kinetochore targeting before other phosphorylations. Previously, it had been demonstrated that HEC1 kinetochore localization was regulated by PP2A-B55, while MIS12 localization was more likely under PP1 control (Gascoigne and Cheeseman, 2013). However, no phosphatase dependency had been established for KNL1. To identify the phosphatase regulating KNL1 kinetochore targeting, we performed live cell imaging of the HeLa cell lines expressing endogenously GFP-tagged KNL1, MIS12 and HEC1 under conditions where PP1α and γ (from here-on referred to as PP1) or PP2A-B55 had been depleted (Figure 4A, Figure S3F). As reported previously, depletion of PP2A-B55 significantly delayed HEC1 disappearance from kinetochores in anaphase but had only a minor effect on KNL1 and MIS12 levels at late stages of anaphase (Figure 4A) (Gascoigne and Cheeseman, 2013). PP1 depletion, however, significantly delayed the release of both MIS12 and KNL1 from kinetochores in early anaphase. For KNL1 specifically, the sharp decline of kinetochores levels observed at the metaphase-to-anaphase transition, was changed into a much more gradual decline over the first 20 minutes of anaphase. Based on this observation we concluded that KNL1 binding to NSL1, and hence recruitment to kinetochores, was regulated by the opposing activities of CDK1-cyclin B1 and PP1. To corroborate this conclusion, PP1 or PP2A-B55 were depleted in the KNL1-GFP HeLa cells, and we asked whether, in the spindle checkpoint re-activation assay, KNL1 localization was maintained in the presence of CDK1 inhibitor when PP1 was absent. Indeed, our data confirmed that in control cells KNL1 was lost from kinetochores when CDK1 was inhibited, but not in cells in which PP1 had been depleted and to a reduced extent when PP2A-B55 was depleted (Figure 4B and C). This confirms that PP1 counteracts CDK1 phosphorylation of NSL1 necessary for KNL1 kinetochore localization. Interestingly, PP1 depletion resulted in an increase of KNL1 kinetochore localization above the control levels, suggesting that PP1 may regulate KNL1 kinetochore localization in more than one way. To test whether the regulation of KNL1 kinetochore levels by the opposing actions of CDK1 and PP1 could explain the loss of kinetochore spindle checkpoint proteins upon CDK1 inhibition, BUB1 localization in cells expressing the GFP-MIS12-MPS1 fusion protein was investigated. As observed before, treatment of mitotically arrested cells with CDKi, even in the presence of kinetochore-tethered MPS1, resulted in significantly reduced BUB1 kinetochore levels (Figure 4D). However, when the cells had been depleted of PP1, but not PP2A-B55, before inhibition of CDK1, BUB1 levels were maintained (Figure 4D and E). This confirms that the interplay between CDK1-cyclin B1 and PP1 indirectly regulates the recruitment of spindle checkpoint proteins through controlling the kinetochore levels of the spindle checkpoint assembly platform KNL1.

**Figure 4.**
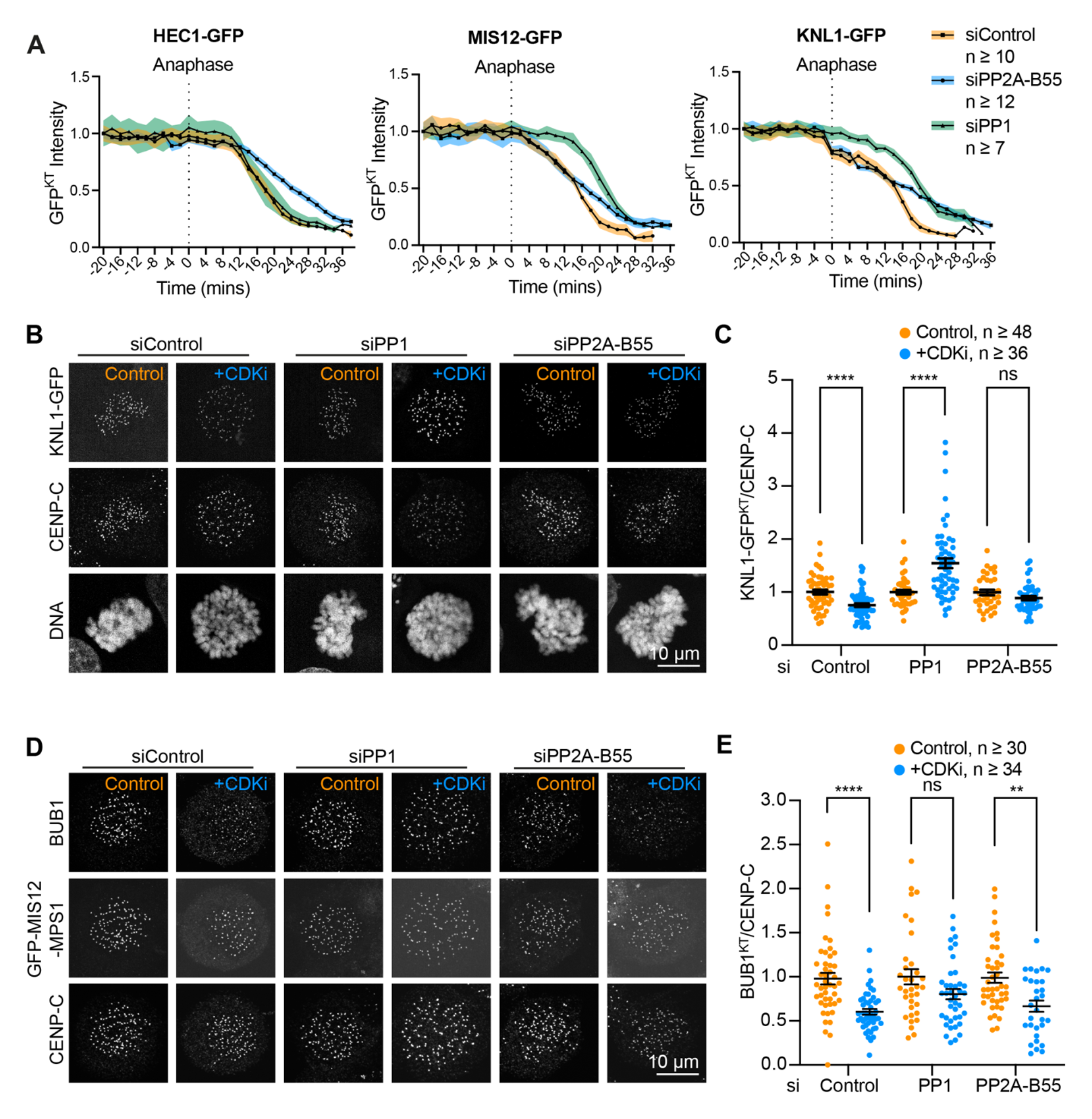
PP1 regulates dissociation of KNL1 from kinetochores. (**A**) HeLa cells expressing HEC1-GFP, MIS12-GFP or KNL1-GFP were depleted of PP1, PP2A-B55 or a control siRNA as indicated for 72 h, then synchronised for 18 h using 2 mM thymidine. Cells were incubated with 100 nM SiR-DNA and imaged 9 h after thymidine release. Graphs show mean cell kinetochore intensities of HEC1-GFP, MIS12-GFP or KNL1-GFP at 2 min intervals as a fraction of the maximum intensity. Shaded regions indicate the SEM. (**B**) HeLa cells expressing KNL1-GFP cells were depleted of PP1, PP2A-B55 or a control siRNA as indicated. After 72 h cells were arrested for 2.5 h with 20 μM MG132, then treated with 3 μM nocodazole for 5 min, fixed and immunostained as indicated. Where indicated, 5 μM flavopiridol was added 1 min before nocodazole treatment (+CDKi). (**C**) Graph shows mean cell intensities of kinetochore KNL1-GFP relative to CENP-C. Bars indicate the SEM. (D) HeLa Flp-In/T-REx GFP-MIS12-MPS1 cells depleted of PP1, PP2A-B55 or a control siRNA as indicated were arrested for 2.5 h with 20 μM MG132, then treated with 3 μM nocodazole for 5 min, fixed and immunostained as indicated. Where indicated, 5 μM flavopiridol was added 1 min before nocodazole treatment (+CDKi). (E) Graph shows mean cell intensities of kinetochore BUB1 relative to GFP-MIS12-MPS1. Bars indicate the SEM.

### NSL1 dephosphorylation by PP1 promotes permanent SAC silencing

To investigate the importance of the CDK1-PP1 regulatory mechanism for the silencing of the spindle assembly checkpoint, HeLa Flp-In T-REx cells stably expressing a GFP-tagged form of the spindle assembly checkpoint marker MAD2 (Hayward et al., 2019) as well as either mCherry-NSL1^WT^ or mCherry-NSL1^2TD^ were filmed progressing through mitosis (Figure 5A-B, Figure S3E). As expected, replacing endogenous NSL1 with mCherry-NSL1^WT^, fully rescued the NSL1 depletion phenotype, and these cells exited mitosis in a timely fashion (Figure 5A, C). Cells expressing GFP-NSL1^2TD^ displayed largely normal mitotic timing (Figure 5C). Interestingly, though, in comparison to their wild type counterparts, cells expressing GFP-NSL1^2TD^ frequently displayed single GFP-MAD2 foci in metaphase and early anaphase (Figure 5B, D). In these cells, spindle assembly checkpoint silencing appeared to have been finalized normally once a metaphase plate had been achieved. However, after SAC silencing had been completed, single GFP-MAD2 foci re-appeared (Figure 5B, arrowheads). These GFP-MAD2 foci did not seem to result in delayed progression through anaphase in line with published reports (Dick and Gerlich, 2013). However, their more frequent appearance in the GFP-NSL1^2TD^ cells indicates that the disassembly of the NSL1-KNL1 interaction via PP1-mediated dephosphorylation of NSL1 is an additional mechanism to shut down spindle checkpoint signalling in early anaphase while maintaining microtubule attachment to kinetochores, hence guaranteeing safe and uni-directional passage through anaphase (Figure 5E).

**Figure 5.**
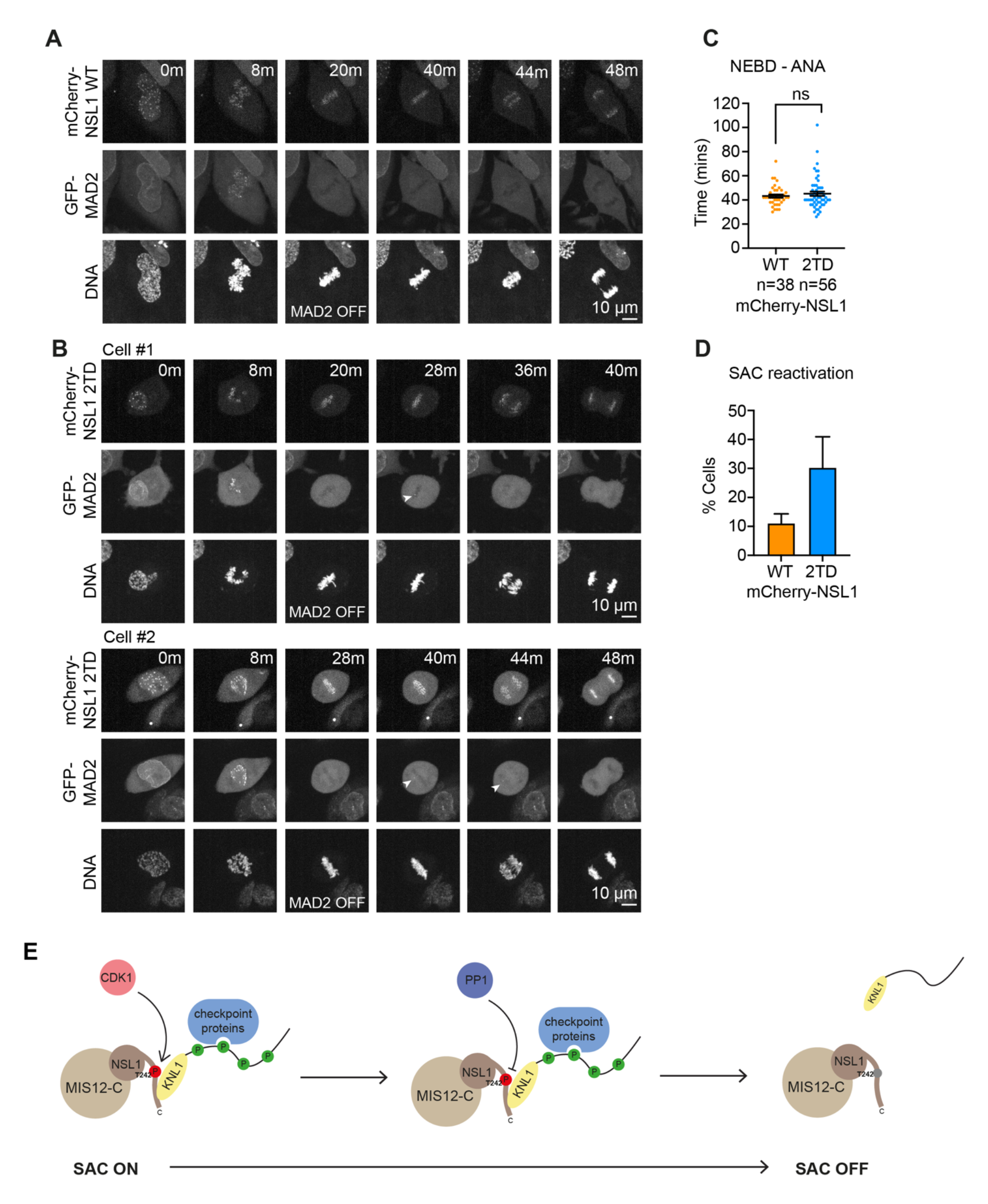
NSL1 dephosphorylation promotes permanent spindle checkpoint silencing. (**A**) Representative stills of HeLa Flp-In/T-REx mCherry-NSL1^WT^ cells, expressing GFP-MAD2 incubated with 100 nM SiR-DNA from NEBD to anaphase. Cells were synchronised for 18 h using 2 mM thymidine and imaged 9 h after thymidine release. (B) Representative stills of HeLa Flp-In/T-REx mCherry-NSL1^2TD^ cells expressing GFP-MAD2 incubated with 100 nM SiR-DNA from NEBD to anaphase. Cells were synchronised for 18 h using 2 mM thymidine and imaged 9 h after thymidine release. Arrowheads indicate MAD2-positive kinetochores in metaphase and anaphase. (C) Graph shows mean time from NEBD to anaphase (mins) for HeLa Flp-In/T-REx mCherry-NSL1^WT^ and mCherry-NSL1^2TD^ cells. Bars indicate SEM. (**D**) Graph shows mean % of cells that showed MAD2-positive kinetochores during metaphase and/or anaphase from a total of 46 mCherry-NSL1^WT^ or 59 mCherry-NSL1^2TD^ cells over 5 or 4 experimental repeats, respectively. Bars indicate SEM. (**E**) Schematic depicting the opposing roles of CDK1 and PP1 in KNL1 kinetochore dissociation to promote timely and irreversible spindle assembly checkpoint silencing.

The metaphase-to-anaphase transition is a critical point in the cell division process, and is therefore safeguarded at several levels. Most importantly, cells will only enter anaphase once spindle assembly checkpoint signalling has been silenced, indicating that all kinetochores have been stably end-on attached to microtubules. SAC silencing involves a number of interconnected processes including the removal of SAC promoting phosphorylations by kinetochore-localized phosphatases, and dynein-mediated stripping and transport of spindle checkpoint components to the spindle poles (Lara-Gonzalez et al., 2021). Once the spindle checkpoint silenced state has been achieved, disassembly of the mitotic checkpoint complex will take over, resulting in activation of the anaphase promoting complex which in turn will lead to activation of separase and ultimately cohesion cleavage. It is paramount that the spindle checkpoint is not re-activated upon loss of sister chromatid cohesion despite the altered tension state across kinetochores. It has been shown previously that the destruction and inactivation of CDK1-cyclin B1 is crucial to prevent SAC reactivation upon anaphase onset (Clijsters et al., 2014; Rattani et al., 2014; Vazquez-Novelle et al., 2014). We propose that prompt removal of KNL1 from the outer kinetochore by dephosphorylation of the CDK1-phosphorylated KNL1 kinetochore binding partner NSL1 contributes to this so-called “point-of-no-return” after which the spindle checkpoint cannot be re-activated anymore, and the cell is committed to anaphase entry (Figure 5E). This differential loss of KNL1 from kinetochores is achieved by the use of PP1 as the dephosphorylating enzyme. In contrast to PP2A-B55, which during mitosis is indirectly inhibited by CDK1 via the Greatwall/ENSA pathway, and is re-activated at anaphase onset with a time delay (Cundell et al., 2013; Cundell et al., 2016; Holder et al., 2019), PP1 is directly inhibited by CDK1-mediated phosphorylation of a conserved, C-terminal threonine (Dohadwala et al., 1994; Kwon et al., 1997), which can be reversed rapidly upon mitotic exit (Bancroft et al., 2020). The differential use of PP2A-B55 and PP1 as opposing phosphatases in the regulation of outer kinetochore proteins hence determines the temporal disassembly characteristics of the different kinetochore components and enables the termination of spindle checkpoint signalling while keeping microtubule-kinetochore attachments intact.

We have previously shown that the dynamic interplay between CDK1-cyclin B1, PP2A-B55 and their substrate MPS1 delineates the spindle checkpoint responsive period during mitosis. The relationship between CDK1-cyclin B1 and PP1 acting on the NSL1-KNL1 interaction that we describe here, reinforces this mechanism by selectively removing the key substrate of MPS1 during SAC signalling, the spindle checkpoint scaffold KNL1. Together, these mechanisms ensure that once CDK1 inactivation has passed a threshold in anaphase, the spindle checkpoint cannot be re-activated anymore.

## Materials and Methods

### Reagents and antibodies

General laboratory reagents were obtained from Sigma-Aldrich and ThermoFisher Scientific unless specifically indicated. Inhibitors were obtained from Tocris Bioscience (PP1 and PP2A inhibitor Calyculin A (1336)); CDK inhibitor Flavopiridol (3094); protein phosphatase inhibitor Okadaic acid (1136)), Santa Cruz Biotechnology (Proteasome inhibitor MG132 (sc-201270)), Millipore (Thymidine (6060-5MG)), and Merck Chemicals Ltd (microtubule polymerization inhibitor Nocodazole (484728-10MG)). Transfection reagents oligofectamine (Invitrogen) for siRNA oligos, and LT1 (Mirus-Bio) for plasmids were used according to the manufacturers’ instructions. Opti-MEM medium (Gibco) was used to set up transfections.

Commercially available antibodies were used for CENP-C (guinea pig pAb; MBL (PD030); 1:2000 dilution for immunofluorescence), BUB1 (rabbit pAb, Bethyl A300- 373A; 1:5000 dilution for immunofluorescence), BUBR1 (rabbit pAb; Bethyl A33-386A; 1:2500 dilution for immunofluorescence), beta Actin (mouse mAb AC-15 HRP conjugate; Abcam (ab49900); 1:5000 dilution for western blotting), HEC1 (mouse mAb 9G3.23; GeneTex (GTX70268); 1:1000 dilution for immunofluorescence, or rabbit pAb; Abcam (ab109496); 1:1000 dilution for western blotting), MAD1 (rabbit pAb; GeneTex (GTX10507); 1:2000 dilution for immunofluorescence), MPS1 (mouse mAb (N1); Abcam (ab11108); 1:2000 dilution for western blotting), NSL1 (rabbit pAb (anti-DC8), Bethyl (A300-795A); 1:1000 dilution for immunofluorescence, 1:1000 dilution for western blotting). Sheep antibodies against mCherry and GFP have been described (Bastos and Barr, 2010). Antibodies against KNL1 and MIS12 were raised in sheep against recombinant GST-KNL1^728-1200^ and recombinant full-length GST-MIS12, respectively, and affinity purified against the same recombinant protein. All secondary antibodies were used at 1:1000 dilutions based on recommended stock concentrations. Secondary donkey antibodies against mouse, rabbit or sheep and labelled with Alexa Fluor 350, Alexa Fluor 555, Alexa Fluor 647 were purchased from ThermoFisher Scientific. Secondary donkey antibodies against guinea pig labelled with Alexa Fluor 647 and secondary donkey antibodies against mouse, rabbit or sheep labelled with HRP were purchased from Jackson ImmunoResearch Laboratories. Protein A and Protein G conjugated to HRP were purchased from Merck. DNA dye Hoechst 33342 was purchased from ThermoFisher Scientific and used at a final concentration of 5 µg/ml.

For western blotting, proteins were separated by SDS-PAGE and transferred to nitrocellulose using a Trans-blot Turbo system (Bio-Rad). Protein concentrations were measured by Bradford assay using Protein Assay Dye Reagent Concentrate (Bio-Rad). All western blots were revealed using ECL (GE Healthcare).

### Molecular biology

Human MPS1 and NSL1 were amplified from Human testis cDNA (Marathon cDNA; Takara Bio Inc.) using Pfu polymerase (Promega). Mammalian expression constructs were made using pcDNA5/FRT/TO (Invitrogen) modified to encode EGFP-, mCherry-or FLAG-reading frames. Generation of the GFP-MIS12-MPS1 fusion construct has been previously described, as has generation of MPS1 kinase dead mutants (Hayward et al., 2019). Mutagenesis was performed using the QuickChange method (Agilent Technologies). DNA primers were obtained from Invitrogen. Gibson assembly was carried out using NEBuilder HiFi DNA Assembly Master Mix (NEB) according to the manufacturer’s instruction for primer design and experimental method.

For the knock down of the catalytic subunits of PPP1CA and PPP1CC, small interfering RNA (siRNA) duplexes 5’-UGGAUUGAUUGUACAGAAAUU-3’ and 5’- GCGGUGAAGUUGAGGCUUAUU-3’ targeting the 3’-UTR of PPP1CA and PPP1CC, respectively, were used for 72 hours. PP2A-regulatory subunit B55 was depleted using a combination of four Smartpools against each isoform (L-004824-00, L-003022- 005, L-019167-00, and L-032298-00 for PPP2R2A, PPP2R2B, PPP2R2C, and PPP2R2D, respectively) for 72 hours. MPS1 was depleted using three oligos against the 3′ UTR (5′-UUGGACUGUUAUACUCUUGAA-3′, 5′-GUGGAUAGCAAGUAUAUU CUA-3′, and 5′-CUUGAAUCCCUGUGGAAAU-3′) for 48 hours. NSL1 was depleted using an siRNA duplex 5’- CAUAGUAGAUAUAGCCACA-3’ against the ORF of NSL1 for 48 hours. Control siRNA was performed against GL2 (luciferase) (5’- CGUACGCGGAAUACUUCGAUU-3’) (Dharmacon #D-001100-01-20).

### Cell culture procedures

HeLa and HEK-293T cells were cultured in DMEM with 1% (vol/vol) GlutaMAX (Life Technologies) containing 10% (vol/vol) bovine calf serum at 37°C and 5% CO_2_. HeLa cells with inducible single integrated copies of a desired transgene were created using the T-REx doxycycline-inducible Flp-In system (Tighe et al., 2008). For cell line generation, 100 ng of pcDNA5/FRT/TO/EGFP or pcDNA5/FRT/TO/mCherry vectors were co-transfected into Flp-In T-REx doxycycline transactivator HeLa cells with 900 ng of the Flp recombinase plasmid pOG44. Cells were selected in 4 μg/ul blasticidin and 200 μg/ul hygromycin B for 2 weeks. Immunofluorescence and western blotting were used to screen for successful, homogenous clones. Clones were then checked in a doxycycline negative background to confirm there was no leaky expression.

Transgene expression was induced by addition of 2 μM doxycycline for a minimum of 24 h.

Heterozygous CRISPR/Cas12a-edited HeLa cells with inserted GFP tags at the C-termini of HEC1, KNL1 or MIS12 were generated following a published PCR tagging method (Fueller et al., 2020). In brief, a gene-specific PCR cassette was generated using two gene specific tagging oligos which provided the homology arms and a generic cassette containing the GFP tag, selection marker and Cas12a-specific crRNA gene (Addgene 120016). PCRs were carried out using HiFi polymerase (Takara Bio UK Ltd) according to the manufacturer’s instructions. The PCR cassette was transfected into HeLa cells with a helper plasmid containing AsCas12a endonuclease (Addgene 89353). Following cleavage, insertion of the PCR cassette into the genome by homologous recombination yielded a fusion of the GFP tag and the gene of interest. Antibiotic resistant clones were selected and successful modification confirmed by immunofluorescence and western blotting. Live cell imaging was used to confirm normal mitotic timing in CRISPR/Cas12a edited cell lines.

Generation of HeLa cells expressing GFP-MAD2 has been previously described (Hayward et al., 2019).

### Transient transfection for protein expression

Transient protein expression for in vitro kinase assays and mass spectrometry was carried out in HEK-293T cells. Cells were seeded 24 h prior to transfection. Transfection mixtures were prepared with 24 μl LT1 (Mirus-Bio), 800 μl Opti-MEM (Gibco) and 8 μg plasmids encoding GFP-or FLAG-tagged constructs. 0.3 μM nocodazole (Merck) was added 30 h after transfection, and cells were incubated for a further 18 h. Cells were harvested by mitotic shake-off and processed depending on the subsequent analysis.

### RNAi rescue assays

RNAi rescues assays with GFP-MIS12-MPS1, GFP-NSL1 or mCherry-NSL1 transgenes were all performed using the same protocol. Transgene induction was initiated with the addition of 2 µM doxycycline (InvivoGen) 3 h prior to siRNA addition. Endogenous MPS1 or NSL1 was depleted for 48 hours. A second induction was performed 24 h into the siRNA depletion. Successful RNAi rescue was confirmed by western blotting for all cell lines.

### Checkpoint response assays

For the “checkpoint re-activation assay” (Hayward et al., 2019; Vleugel et al., 2015) cells were treated with 20 μM MG132 for 2.5 h. At 5 min before fixation, 3 μM nocodazole was added. If CDK1 was inhibited, 5 μM flavopiridol was added 1 min or 5 min before nocodazole treatment, with DMSO at the same volume used as a control. For the “checkpoint maintenance assay” cells were treated with 0.3 μM nocodazole for 2 h. At 30 min before fixation, 20 μM MG132 was added to prevent mitotic exit. If CDK1 was inhibited, 5 μM flavopiridol was added 6 min before fixation, with DMSO at the same volume used as a control. All cells were fixed in PTEMF and immunostained as indicated in figure legends before imaging.

### Immunofluorescence staining

Cells were fixed with PTEMF buffer (20 mM PIPES-KOH pH 6.8, 0.2% (vol/vol) Triton X-100, 10 mM EDTA, 1 mM MgCl_2_, 4% (vol/vol) formaldehyde) for 12 min at room temperature. Coverslips were washed in phosphate buffered saline (PBS) and incubated in blocking buffer (3% (wt/vol) bovine serum albumin in PBS) for a minimum of 45 min. Coverslips were incubated face-down on 80 µl droplets of primary antibodies in a humidified chamber for 1.5 h. Following primary antibody incubation coverslips were washed 3x in PBS. Secondary donkey antibodies against mouse, rabbit, guinea pig, or sheep, labelled with Alexa Fluor 405, Alexa Fluor 555, or Alexa Fluor 647 (Molecular Probes) were used at 1:1000. Coverslips were incubated face-down on 80 µl droplets of diluted antibodies in a humidified chamber for 45 min. Coverslips were washed 3x in PBS and 1x in distilled deionised water. Coverslips were left to dry completely before being mounted with 6 µl of Mowiol 4-88 (Sigma) according to manufacturer’s instructions.

### Immunofluorescence microscopy

Imaging was performed on a DeltaVision Core light microscopy system (GE Healthcare) using a 60x/1.35 NA objective fitted to an Olympus IX-71 microscope stand. Standard filter sets for DAPI, FITC and Cy5, a CoolSnap HQ2 charge-coupled device (CCD) camera (Photometrics) and the software package softWoRx (GE Healthcare) were used. Cells were imaged using a 0.2 μm interval and a total stack of 4 μm, and deconvolved for presentation using softWoRx.

For quantitation, imaging was performed using a 60x/1.35 NA oil immersion objective on a BX61 Olympus microscope with filter sets for DAPI, EGFP/Alexa Fluor 488, Alexa Fluor 555, Alexa Fluor 647 (Chroma Technology Corp.) to sequentially excite and collect fluorescence images on a CoolSNAP HQ2 camera (Roper Scientific) using MetaMorph 7.5 imaging software (GE Healthcare). Cells were imaged using a 0.2 μm interval and a total stack of 2.4 μm.

### SoRa Confocal microscopy

Super-resolution immunofluorescence microscopy was performed on an Olympus SoRa spinning disk confocal microscope using a 60x/1.5 NA objective fitted to an Olympus IX-83 microscope stand with 3.2x optical zoom and a Yokogawa CSU-W1 SoRa super-resolution spinning disk. Solid state lasers emitting 405 nm, 488 nm, 561 nm and 633 nm were used. Images were captured with a Prime BSI sCMOS camera (photometrics) using the Olympus cellSens software package. Images were acquired with a 0.2 µm interval over a total distance of 4 µm. Super-resolution enhancement and constrained iterative deconvolution was performed on cellSens.

### Live cell imaging

For time lapse imaging, cells were seeded on circular glass bottom Fluorodish imaging dishes (World Precision Instruments). Cells were treated for 18 h with 2 mM thymidine. The thymidine was removed by washing 6x with PBS. After washes, the PBS was replaced with Fluorobrite media (ThermoFisher Scientific) supplemented with 10% (vol/vol) bovine calf serum and 1% (vol/vol) GlutaMAX (Life Technologies). SiR-DNA (Spirochrome) was added at a concentration of 100 nM. Imaging commenced 9 h after thymidine release at 37°C, 5% CO_2_ on an Olympus SoRa spinning disk confocal microscope using a 60x/1.5 NA objective fitted to an Olympus IX-83 microscope stand with a Yokogawa CSU-W1 SoRa super-resolution spinning disk. Solid state lasers emitting 488 nm, 561 nm and 633 nm were used. Images were captured with a Prime 95B sCMOS camera (photometrics) using the Olympus cellSens software package. Images were acquired with a stack of 12 z-planes spaced approximately 1 µm apart, every 2 min for 5 h.

### Image processing and analysis

Image processing and analysis was performed using the FIJI distribution of ImageJ (Schindelin et al., 2012).

For figures, images acquired on a BX61 Olympus microscopy system or deconvolved images acquired on either a DeltaVision Core light microscopy system or an Olympus SoRa spinning disk confocal microscopy system were maximum projected and saved as 8-bit TIFF files. TIFF files were imported into Illustrator CS6 (Adobe) for figure production. Quantitation was performed on images of cells from a BX61 Olympus microscopy system which were cropped to 250 x 250 px and sum projected through 7 z-slices. Kinetochore intensities were determined by placing 8 px-diameter circular ROIs at the maxima of kinetochores and measuring mean pixel intensity of each channel within selections. Where possible, 20 kinetochores were measured per cell. Background measurements were generated by taking an equivalent number of pixels as were in the ROI as close to the ROI without overlapping with kinetochores. Data analysis was performed in Excel (Microsoft). Kinetochore signal intensities were background-adjusted by subtracting the background signal on a channel-by-channel basis. The mean intensity of the channel of interest was then divided by the mean intensity of either the CENP-C or GFP channel as stated on a per-kinetochore basis. Mean kinetochore localisation intensities were then calculated for each cell. Normalisation was performed within repeats by dividing the cell’s mean kinetochore localisation intensity by that of the group which was being normalised to.

For measurement of mitotic timings, maximum-projected images of time lapse imaging data from an Olympus SoRa spinning disk confocal microscopy system were used. The times at which NEBD, metaphase, and anaphase occurred were quantified based on DNA morphology. For measurement of kinetochore intensity from time lapse imaging, maximum-projected images of time lapse imaging data from an Olympus SoRa spinning disk confocal microscopy system were used. 10x 8 px-diameter circular ROIs were placed at the maxima of GFP positive kinetochores and mean pixel intensity at the first timepoint they were visible was measured. The same ROIs were then measured at previous timepoints. 10x 8 px-diameter circular ROIs at the maxima of GFP positive kinetochores were then measured for each subsequent timepoint until they were no longer visible. The same ROIs as the final visible timepoint were then measured at subsequent timepoints. Background measurements were taken from 10 chromatin free cytoplasmic regions. For figures, individual cells from specific timepoints were cropped and saved as 8-bit TIFF files. TIFF files were imported into Illustrator CS6 (Adobe) for figure production.

All statistical analysis and graph production was performed in GraphPad Prism version 9.2.0 (GraphPad Software). Unless otherwise stated, each cell measured was considered as a biological replicate (n), hence the mean cell averages of kinetochores was used for statistical analysis. At least three independent repeats of each experiment were performed, with statistical analysis performed using a sum of biological replicates from all independent experiments. Data sets were tested for normal distribution via a D’Agostino-Pearson omnibus K2 test. If all groups in an experiment were normally distributed, then the means were compared using an unpaired 2 tailed t-test (with Welch’s correction if the groups had unequal standard deviations). If all groups did not exhibit a normal distribution, then medians were compared using a Mann-Whitney test. Unless otherwise stated, graphs display the mean ±SEM. P-values are shown on graphs as follows: p > 0.05 = not significant (ns), p ≤ 0.05 = *, p ≤ 0.01 = **, p ≤ 0.001 = ***, p ≤ 0.0001 = ****.

### Analysis of NSL1 phosphorylation

HeLa cells were arrested in 0.3 µM nocodazole for 18 h and harvested by mitotic shake-off. For asynchronous cell population, HeLa cells were harvested using 1 mM EDTA. Cell pellets were lysed and subsequently incubated with λ-protein phosphatase (NEB) for 1 h at 30°C according to the manufacturer’s instructions. 1x Laemmli buffer was added and samples denatured at 100°C for 5 min. Samples were run on PhosTag or SDS-PAGE gels and analysed by western blotting. For PhosTag gels, 10% (wt/vol) polyacrylamide separating gels were prepared with 100 µM MnCl_2_ and 50 µM PhosTag reagent (Wako Chemicals, AAL-107S1). Typically, 10 µg of lysate was loaded for PhosTag gels. Prior to transfer, gels were equilibrated with 3x 3 min washes in transfer buffer (20 mM Tris-HCl; 150 mM Glycine, 0.1% (wt/vol) SDS, 20% (vol/vol) MeOH, 20 mM EDTA) and subsequently 3x 3 min washes in transfer buffer without EDTA (20 mM Tris-HCl, 150 mM Glycine, 0.1% (wt/vol) SDS, 20% (vol/vol) MeOH).

### Immunoprecipitation assays

Cells were grown in 15 cm dishes (number of dishes varied depending on the size of immunoprecipitation). For GFP-NSL1 Flp-In cell lines, transgene expression was induced by addition of 2 μM doxycycline for 24 h. In all experiments, cells were arrested in 0.3 µM nocodazole for 18 h and harvested by mitotic shake-off. Cells were pelleted at 500 xg for 4 min at room temperature, washed with PBS and pelleted again. If CDK1 was inhibited, pellets were resuspended in 1 ml Opti-MEM and 5 μM flavopiridol was added for 10 min, with DMSO at the same volume used as a control. Further dephosphorylation was stopped after 10 min by addition of 25 nM Calyculin A and cells were pelleted at 500 xg for 4 min at room temperature.

Pellets were lysed in lysis buffer (20 mM Tris-HCl pH 7.4, 150 mM NaCl, 1% (vol/vol) IGEPAL, 0.1% (wt/vol) sodium deoxycholate, 100 nM okadaic acid, 40 mM sodium β- glycerophosphate, 10 mM NaF, 0.3 mM sodium vanadate, 1:250 protease inhibitor cocktail (Sigma), 1:100 phosphatase inhibitor cocktail (Sigma)) for 30 min at 4°C before being clarified at 14,000 rpm for 15 min at 4°C. Per 1 mg of lysate, 1 µg appropriate antibody was added (homemade sheep anti-GFP or homemade sheep anti-mCherry as a negative IgG control). Anti-GFP immunoprecipitations were performed using Protein-G Dynabeads (Invitrogen), anti-FLAG immunoprecipitations were performed using anti-FLAG M2 magnetic beads (Merck). All beads were washed 3x in lysis buffer prior to use. Immunoprecipitations were performed at 4°C for 2 h. Beads were washed 6x, 3x in lysis buffer and 3x in wash buffer (20 mM Tris-HCl pH 7.4, 150 mM NaCl, 0.1% (vol/vol) IGEPAL, 40 mM sodium β-glycerophosphate, 10 mM NaF, 0.3 mM sodium vanadate). Beads were separated from the flow-through using a magnet. If samples were used for western blot analysis, 1x Laemmli buffer was added and samples denatured at 100°C for 5 min.

### MPS1 kinase assays

For kinase assays, 1 µg recombinant FLAG-MPS1 WT/KD/S281A/4A purified by immunoprecipitation on anti-FLAG M2 magnetic beads from HEK-293T cells was auto-phosphorylated for 10 min at 30°C in 50 mM Tris-HCl pH 7.3, 50 mM KCl, 10 mM MgCl_2_, 20 mM sodium β-glycerophosphate, 15 mM EGTA, 0.1 mM ATP, 1 mM DTT, and 1 µCi [^32^P]γ-ATP per reaction. The reaction was terminated by addition of 2x Laemmli buffer and samples run on SDS-PAGE gels. Incorporation of γ^32^P into FLAG-MPS1 was used as a readout of kinase activity.

### Mass spectrometry

GFP-NSL1 was immunoprecipitated from HEK-293T cells that had been transiently transfected with GFP-NSL1. Cell harvesting and immunoprecipitation was carried out as previously described, with the only variation of reduced IGEPAL in buffers: 0.5% (vol/vol) IGEPAL in lysis buffer and no IGEPAL in wash buffer. Following immunoprecipitation, Protein-G Dynabeads (Invitrogen) were washed 3x in PBS at 4°C.

Samples were digested using lysyl endopeptidase (R) and trypsin. Peptides were separated by nano liquid chromatography (ThermoFisher Scientific Easy-nLC 1000) coupled in line to a Q Exactive mass spectrometer equipped with an Easy-Spray source (ThermoFisher Scientific). Peptides were trapped onto a C18 PepMac100 precolumn (300 µm i.d.x5 mm, 100 Å, ThermoFisher Scientific) using Solvent A (0.1% Formic acid, HPLC grade water). The peptides were further separated onto an Easy-Spray RSLC C18 column (75 µm i.d., 50 cm length, ThermoFisher Scientific) using a 30 min linear gradient (15% to 35% solvent B (0.1% formic acid in acetonitrile)) at a flow rate 200 nl/min. The raw data were acquired on the mass spectrometer in a data-dependent acquisition mode (DDA). Full-scan MS spectra were acquired in the Orbitrap (Scan range 350-1500 m/z, resolution 70,000; AGC target, 3e6, maximum injection time, 100 ms). The 10 most intense peaks were selected for higher-energy collision dissociation (HCD) fragmentation at 30% of normalised collision energy. HCD spectra were acquired in the Orbitrap at resolution 17,500, AGC target 5e4, maximum injection time 120 ms with fixed mass at 180 m/z. Charge exclusion was selected for unassigned and 1+ ions. The dynamic exclusion was set to 5 s.

The supplemental material shows the analysis of the effect of MPS1 CDK1 phosphorylation on MPS1 activity (Figure S1), the characterisation of KNL1-GFP, HEC1-GFP and MIS12-GFP CRISPR/Cas12a-edited HeLa cells and live imaging stills of KNL1-GFP, HEC1-GFP and MIS12-GFP (Figure S2), and the analysis of the effect of NSL1 depletion on KNL1, and western blots demonstrating the efficiency of the NSL1 RNAi rescue strategy and the phosphatase RNAi (Figure S3).

## Acknowledgements

ILM was the recipient of a Wellcome Trust PhD studentship. The work was supported by a Cancer Research UK Discovery Programme grant DRCNPG-Nov21\100004 to UG. Mass spectrometry sample preparation and analysis was carried out by Oxford Advanced Proteomics Facility. We thank Francis Barr for critical reading of the manuscript. We acknowledge that this work would not have been possible without the HeLa cell line, which was developed from Henrietta Lacks’ cells taken without compensation or informed consent.

The authors declare no competing financial interests.

For reagent requests, please contact Ulrike Gruneberg.

## Author contributions

Conceptualization: U.Gruneberg. Investigation: I. Lim-Manley. Funding acquisition: I. Lim-Manley; U. Gruneberg. Supervision: U. Gruneberg. Writing – original draft: U. Gruneberg; I. Lim-Manley.

