## Supplemental material for "Spindle checkpoint signalling in anaphase is prevented by KNL1 release from kinetochores"

**Supplemental material**

**Figure S1. Analysis of the effect of CDK1 phosphorylation on MPS1 activity.** (**A**) Schematic of domain organization and reported CDK1 sites of MPS1. (**B**) Western blot confirming successful RNAi rescue with GFP-MIS12-MPS1 fusion protein with 4A/4E mutations. HeLa Flp-In/T-REx GFP-MIS12-MPS1 cells were depleted of endogenous MPS1 and GFP-MIS12-MPS1 transgenes were induced by addition of doxycycline as indicated. * indicates non-specific band. (**C**) HeLa Flp-In/T-REx GFP-MIS12-MPS1 cells depleted of endogenous MPS1 were arrested for 2.5 h with 20 μM MG132, then treated with 3 μM nocodazole for 5 min, fixed and immunostained as indicated. Where indicated, 5 μM flavopiridol was added 1 min before nocodazole treatment (+CDKi). (**D**) Graph shows mean cell intensities of kinetochore BUB1 relative to GFP-MIS12-MPS1. Bars indicate the SEM. (**E**) Western blot analysis of a 10 min in vitro radioactive kinase assay using purified FLAG-MPS1 and [$^{32}$P]gamma-ATP. Wild type (WT), Kinase-dead (KD). (**F**) Graph shows mean $^{32}$P signal relative to amount of FLAG-MPS1. Bars indicate the S.D.

**Figure S2. Characterisation of GFP-kinetochore protein cell lines.** (**A**) Western blot confirming heterozygous clones of KNL1-GFP, MIS12-GFP and HEC1-GFP. Arrowheads indicate GFP-tagged proteins. * indicates non-specific bands. (**B**) Live cell imaging was used to confirm normal mitotic timing in CRISPR/Cas12a edited cells. Graph shows mean time from NEBD to anaphase (mins) of the different cell lines. All CRISPR/Cas12a-tagged cell lines showed mitotic timing very similar to the parental cell line. Bars indicate the SEM. (**C**) Representative movie stills of HeLa cells expressing KNL1-GFP, MIS12-GFP or HEC1-GFP and incubated with 100 nM SiR-

DNA. Cells were synchronised for 18 h using 2 mM thymidine. Cells were imaged 9 h after thymidine release.

**Figure S3. Analysis of NSL1 depletion phenotype.** (**A**) Western blot showing depletion of NSL1 following a 48 h RNAi in HeLa cells. GL2 was used as a control RNAi. (**B**) HeLa cells were control depleted or depleted of endogenous NSL1. After 48 h, cells were fixed and immunostained as indicated. (**C**) Graph shows mean cell intensities of kinetochore BUB1, MAD1, KNL1 or HEC1 relative to CENP-C. Bars indicate SEM. (**D**) Western blot confirming successful RNAi rescue with GFP-NSL1 mutants. HeLa Flp-In/T-REx GFP-NSL1 cells were depleted of endogenous NSL1, and GFP-NSL1 transgenes were induced by addition of doxycycline as indicated. * indicates GFP-NSL1 transgene splice variant or degradation product. (**E**) Western blot confirming successful RNAi rescue with mCherry-NSL1 mutants. HeLa Flp-In/T-REx mCherry-NSL1 cells, also constitutively expressing GFP-MAD2, were depleted of endogenous NSL1, and mCherry-NSL1 transgenes were induced by addition of doxycycline as indicated. (**F**) Western blot showing depletion of PPP1CA and PPP1CC (siPP1) or PPP2R2A (siPP2A-B55) following a 72 h RNAi in HeLa cells. GL2 was used as a control RNAi. * indicates non-specific bands.

#### Figure S1

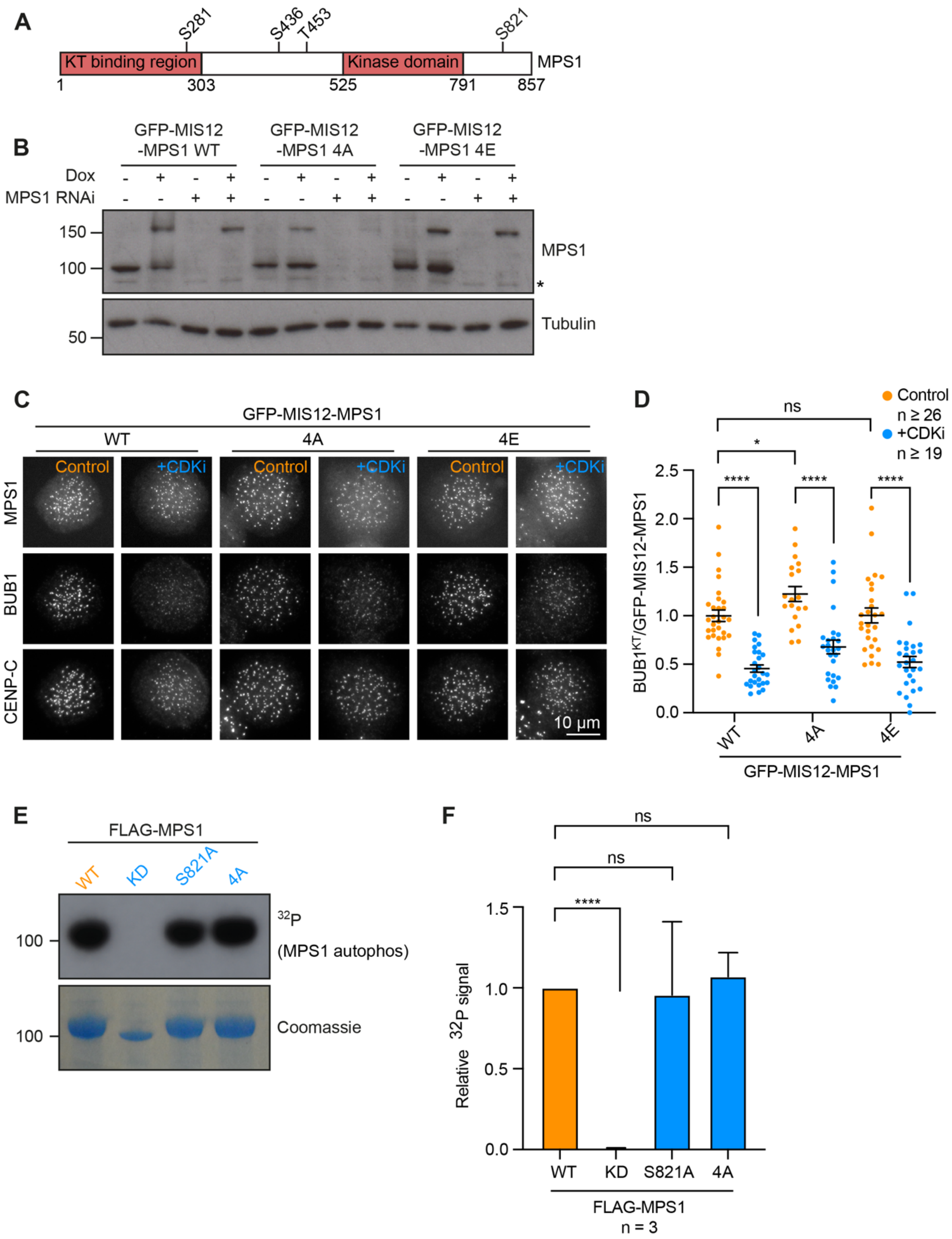

A
S281
S436
T453
S821
KT binding region
Kinase domain
MPS1
1
303
525
791
857
B
GFP-MIS12 -MPS1 WT
GFP-MIS12 -MPS1 4A
GFP-MIS12 -MPS1 4E
Dox
MPS1 RNAi
150
100
50
MPS1
*
Tubulin
C
GFP-MIS12-MPS1
WT
4A
4E
Control
+CDKi
GFP-MIS12-MPS1
BUB1
CENP-C
10 μm
D
Control n ≥ 26
+CDKi n ≥ 19
ns
*
****
BUB1KT/GFP-MIS12-MPS1
2.5
2.0
1.5
1.0
0.5
0.0
WT
4A
4E
GFP-MIS12-MPS1
E
FLAG-MPS1
WT
KD
S821A
4A
100
32P (MPS1 autophos)
Coomassie
F
ns
****
1.5
1.0
0.5
0.0
Relative 32P signal
WT
KD
S821A
4A
FLAG-MPS1
n = 3

#### Figure S2

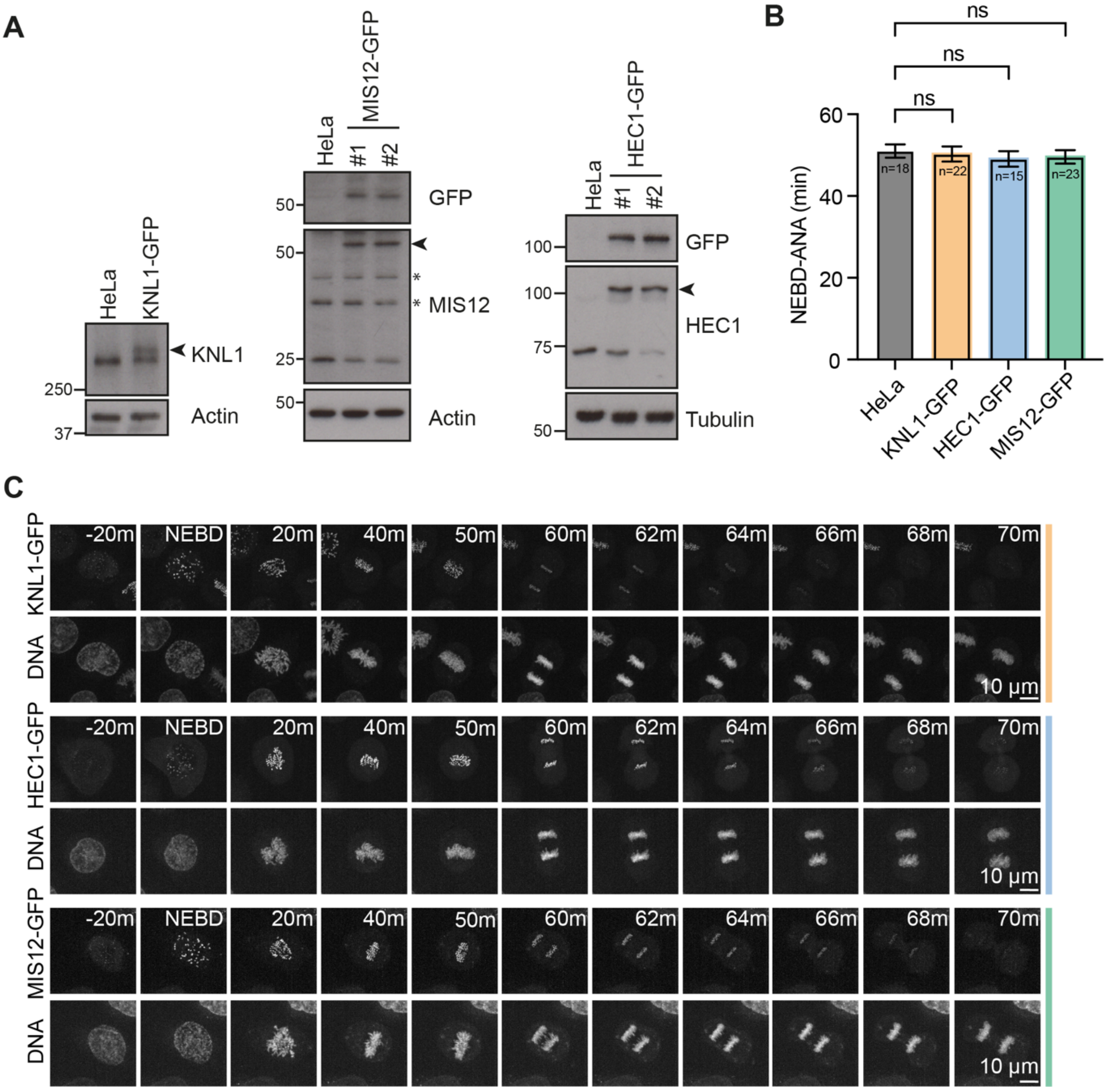
A
HeLa
KNL1-GFP
KNL1
250
37
Actin
MIS12-GFP
#1
#2
GFP
50
MIS12
25
*
HEC1-GFP
HEC1
100
75
Tubulin
B
ns
ns
ns
NEBD-ANA (min)
60
40
20
0
n=18
n=22
n=15
n=23
HeLa
KNL1-GFP
HEC1-GFP
MIS12-GFP
C
DNA
-20m
NEBD
20m
40m
50m
60m
62m
64m
66m
68m
70m
10 μm

### Figure S3

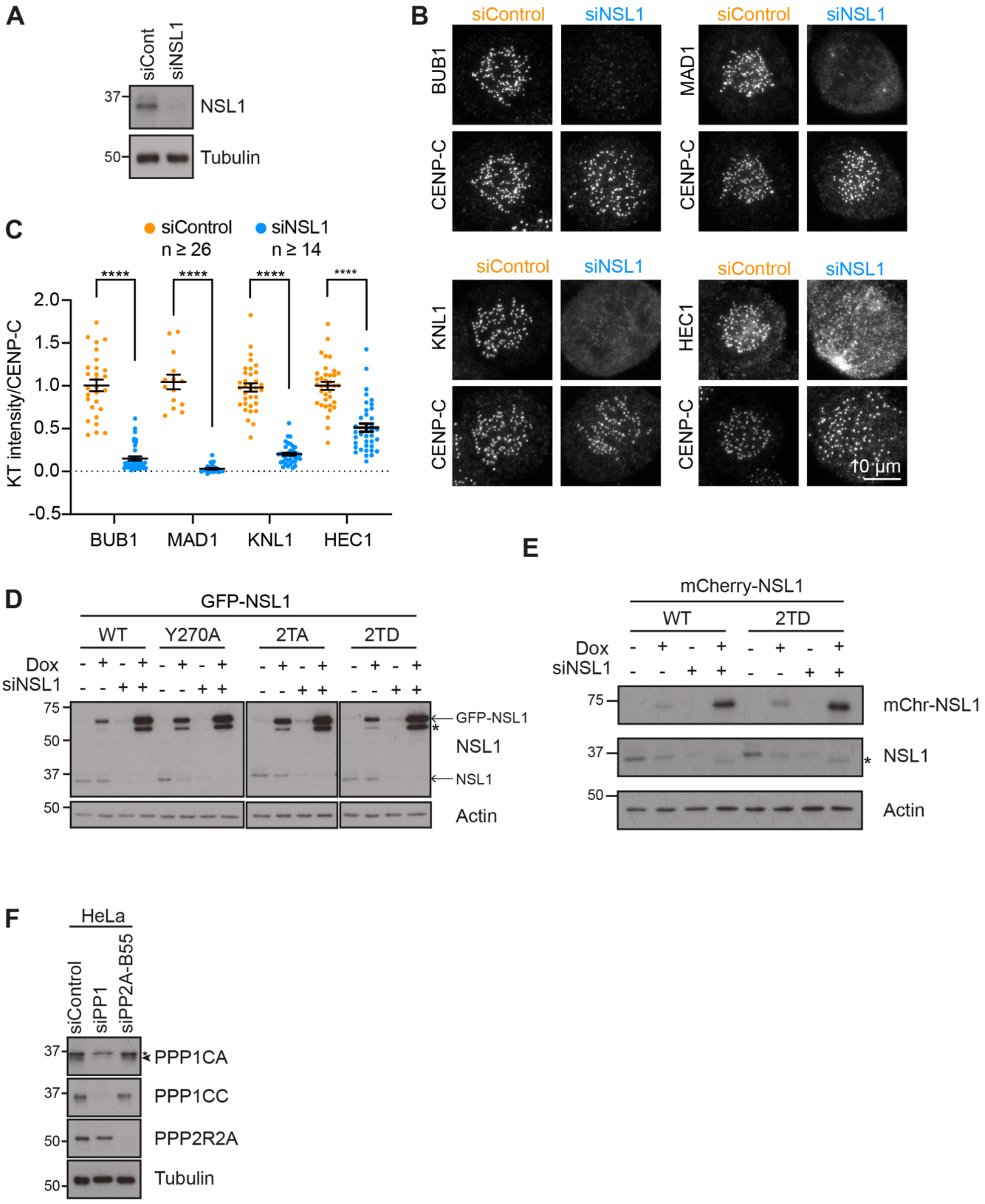
A
siCont
siNSL1
37
NSL1
50
Tubulin
B
siControl
siNSL1
BUB1
MAD1
CENP-C
KNL1
HEC1
10 µm
C
siControl n ≥ 26
siNSL1 n ≥ 14
****
KT intensity/CENP-C
2.0
1.5
1.0
0.5
0.0
-0.5
BUB1
MAD1
KNL1
HEC1
D
GFP-NSL1
WT
Y270A
2TA
2TD
Dox
siNSL1
75
50
37
GFP-NSL1
NSL1
Actin
E
mCherry-NSL1
WT
2TD
Dox
siNSL1
mChr-NSL1
NSL1
Actin
F
HeLa
siControl
siPP1
siPP2A-B55
PPP1CA
PPP1CC
PPP2R2A
Tubulin
